# Decoding mRNA translatability and stability from 5’UTR

**DOI:** 10.1101/2020.03.13.990887

**Authors:** Longfei Jia, Yuanhui Mao, Quanquan Ji, Devin Dersh, Jonathan W. Yewdell, Shu-Bing Qian

## Abstract

Precise control of protein synthesis by engineering sequence elements in 5’ untranslated region (5’UTR) remains a fundamental challenge. To accelerate our understanding of *cis*-regulatory code embedded in 5’UTR, we devised massively parallel reporter assays from a synthetic mRNA library composed of over one million 5’UTR variants. A completely randomized 10-nucleotide sequence preceding an upstream open reading frame (uORF) and downstream GFP leads to a broad range of mRNA translatability and stability in mammalian cells. While efficient translation protects mRNA from degradation, uORF translation triggers mRNA decay in a UPF1-dependent manner. We also identified translational inhibitory elements in 5’UTR with G-quadruplex as a mark for mRNA decay in the P-body. Unexpectedly, an unstructured A-rich element in 5’UTR, while enabling cap-independent translation, destabilizes mRNAs in the absence of translation. Our results not only expose diverse sequence features of 5’UTR in controlling mRNA translatability, but also reveal ribosome-dependent and -independent mRNA surveillance pathways.

## Introduction

In eukaryotic cells, rates of individual mRNA translation and degradation are highly regulated, spanning several orders of magnitude. Recent experimental evidence indicates a complex relationship between these rates across the transcriptome ^1^. While efficient translation typically protects mRNA from degradation ^2^, mRNA surveillance pathways rely on translation to degrade faulty mRNAs ^3^. Emerging evidence suggests that the codon optimality is another determinant of mRNA stability ^4^, implying that ribosome pausing favors mRNA decay. Despite the crucial role of ribosome dynamics in mRNA quality control, whether translation initiation affects the mRNA half-life is uncertain ^5^.

5’ UTR contains key elements of translational regulation, such as structural motifs and upstream open reading frames (uORFs). By controlling the selection of translation initiation sites (TISs), many sequence elements in 5’UTR contribute to mRNA translatability ^6^. Using parallel reporter assays, prior studies have attempted to address the regulatory code of translation initiation ^7–10^. However, it is difficult to dissect the contribution of uORF translation from alternative TISs, which often compete with the main start codon by sequestering ribosomes. Further, the potential effect of 5’UTR variation on mRNA stability has not been considered. Systematic characterization of the *cis*-regulatory elements in 5’UTR will help establish the logical and mechanistic relationships between translation initiation and mRNA decay, if indeed such relationships exist.

Here, we devised massively parallel reporter assays from a synthetic mRNA library to address this fundamental issue. We found that randomized 10-nucleotide sequence preceding a uORF and downstream GFP leads to a broad range of mRNA translatability in mammalian cells. Unexpectedly, variation of a few nucleotides in 5’UTR alters mRNA stability in an order of magnitude. Our results not only expose diverse sequence features of 5’UTR in controlling mRNA translatability, but also reveal ribosome-dependent and -independent mRNA surveillance pathways initiated from 5’UTR.

## Results

We devised a high-throughput uORF reporter assay compatible with fluorescence-activated cell sorting and polysome profiling (Fig. 1a). The reporter harbors a 5’ leader sequence from β-globin followed by GFP with an optimal start codon AUG. We inserted into 5’UTR a sequence encoding SIINFEKL, a tracer peptide that is presented on the cell surface by the mouse major histocompatibility complex class I molecules H-2K^b 11^, and can be detected by the 25D1 monoclonal antibody with exquisite sensitivity (~100 complexes/per cell) ^12^. We replaced the start codon of SIINFEKL with a random 10-nt-long sequence followed by a sequence encoding 5 amino acids from the natural SIINFEKL flanking sequence. These additional amino acids are removed at high efficiency by cellular proteases (Supplementary Fig. 1a). Upon transfection into HEK293 cells stably expressing H-2K^b^, both the tracer peptide and GFP levels can be precisely determined via flow cytometry (Fig. 1a).

**Fig. 1.**
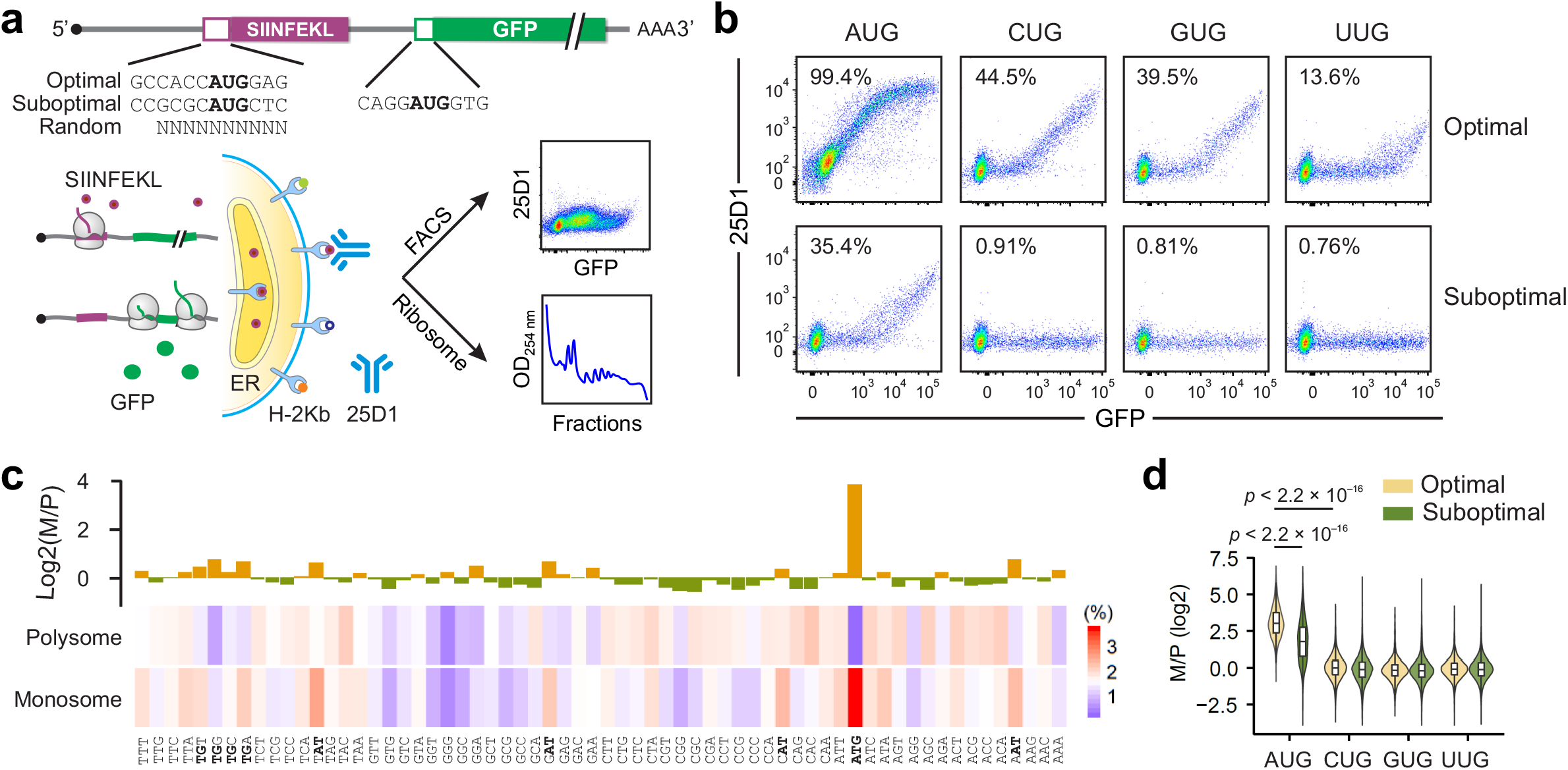
Characterization of start codon selection using synthetic uORF reporters. **(a)** Schematic of uORF reporter assay suitable for FACS and polysome fractionation. The uORF reporter contains a sequence encoding SIINFEKL (purple) followed by GFP (green). The optimal, suboptimal, and random 10-nt sequences are placed before the uORF, whose translation in transfected HEK293-K^b^ cells can be monitored by either flow cytometry or ribosome fractionation. **(b)** Representative flow cytometry scatterplots of HEK293-K^b^ cells transfected with plasmids of uORF reporters with optimal context (top panel) or suboptimal context (bottom panel) of AUG, CUG, GUG or UUG codons. Values shows the percentage of 25D1^+^ population over the total. **(c)** HEK293-K^b^ cells were transfected with mRNA reporters followed by ribosome fractionation. A bar plot (top) shows the ratio of triplet frequency within the random sequences enriched in monosome and polysome. The original frequency of triplets in different ribosome fractions is shown as a heat map (bottom). The triplets ATG, NAT and TGN are highlighted. **(d)** A violin plot shows the ratio of relative abundance of AUG-like triplets between monosome and polysome (M/P). The optimal sequence context has a purine at position −3 and a guanine at position +4. Boxplots show the lower, upper quartile and the median of the M/P ratio. P values are calculated by Wilcox test.

To validate this uORF reporter system, we first examined the efficiency of start codons with different sequence contexts. For an AUG initiator, the initiation efficiency is governed by the adjacent bases with an optimal context having a purine at position −3 and a guanine at position +4. Indeed, the presence of an optimal TIS favored uORF translation at the expense of GFP (Fig. 1b). Altering the sequence context flanking the AUG reduced the uORF-encoded 25D1 signals with a corresponding increase of GFP levels. The reciprocal relationship between SIINFEKL and GFP translation is consistent with the leaky scanning model. We next interrogated uORF translation from non-AUG start codons. In agreement with previous studies ^13–15^, we observed reduced 25D1 signals in a clear order of AUG > CUG > GUG > UUG when the sequence context is optimal (Fig. 1b, top panel). However, when the sequence context is suboptimal, only the AUG triplet enables uORF translation (Fig. 1b, bottom panel).

Notably, the plasmid-based reporter is biased towards GFP expression due to the shorter half-life of K^b^-SIINFEKL complexes on the cell surface than that of intracellular GFP (hours vs. days). To circumvent this problem, we used RNA transfection that allows measurement of translational products accumulated over a 5 hr window. The mRNA reporters increased the 25D1/GFP ratio by approximately 10-fold in transfected cells, as exemplified by the one harboring optimal or suboptimal uTIS codons (Supplementary Fig. 1b – 1c).

To generate a RNA-based reporter library, we designed a PCR-amplification approach using degenerate primers composed of random 10-nt-long sequences that are placed 5’ to the SIINFEKL flanking sequences (Supplementary Fig. 2a). To ensure randomness, we sequenced and compared nucleotide oligos synthesized from different vendors (Supplementary Fig. 2b). With over one million possible sequences (4^10^), we used the pool of PCR products as templates for *in vitro* RNA synthesis followed by 5’ capping and 3’ polyadenylation. Unlike a plasmid-based library, the mRNA library ensures uniformity of transcript variants by excluding possibilities of cryptic promoters, differential transcription, alternative splicing events, and unexpected internal modifications.

Compared to a reporter containing GFP alone, transfection with the pooled uORF reporters resulted in a modest reduction of GFP expression and a slight increase of 25D1 staining (Supplementary Fig. 2c). This is expected, as the majority of random sequences do not specify start codons. We sorted transfected cells into 25D1^H^ and GFP^H^ populations followed by deep sequencing of inserts. Since a single cell could be transfected with multiple mRNA molecules with different sequence variants, individual codons are not segregated well in distinct cell populations (Supplementary Fig. 2d). Nevertheless, many AUG-like codons are enriched in the 25D1^H^ population, whereas GC-rich triplets are over-represented in the GFP^H^ population (Supplementary Fig. 2e). The modest enrichment of AUG-like codons in the 25D1^H^ population suggests that leaky scanning occurs frequently at these TIS sites, resulting in robust signals of both 25D1 and GFP.

To better separate mRNAs with uORF translation only, we collected mRNAs based on the number of associated ribosomes using sucrose gradient. Based on the size of the putative uORFs, only one ribosome can be accommodated at a time. Therefore, mRNAs with active uORF translation are expected to reside in the monosome fraction. By contrast, mRNAs with downstream GFP translation are likely engaged with multiple ribosomes. Indeed, mRNAs uncovered from the monosome show a prominent enrichment of an AUG codon within the insert, which conversely, is highly depleted from polysome-derived mRNAs (Fig. 1c). N**AU** and **UG**N triplets are also overrepresented in monosome mRNAs, another indication of AUG codons with varied flanking sequences.

To examine the sequence context of AUG, we scored the monosome/polysome (M/P) ratio (log2) for sequences with all permutations of NNNNAUGNNN (Supplementary Fig. 3a). A direct comparison of high and low M/P ratio revealed the importance of a purine (A or G) at −3 position and a G at +4 position (Supplementary Fig. 3b). To validate the above sequencing results, we chose several top hits from discrete ribosome fractions and examined their translational status by flow cytometry (Supplementary Fig. 3c). Indeed, monosome-enriched variants (M1 and M2) showed strong 25D1 signals, whereas polysome residents (P1 and P2) expressed GFP only. Notably, non-AUG codons, including near cognate codons, are poorly enriched in the monosome-associated mRNAs, regardless of the sequence context (Fig. 1d). Therefore, the codon identify is more important than the sequence context in controlling leaky scanning.

Although codons enriched in the GFP^H^ population and polysome are positively correlated (*R* = 0.49, Supplementary Fig. 4a), surprisingly, the monosome-enriched AUG codon was poorly recovered from the 25D1^H^ population. One possible explanation is that mRNA stability is inversely proportional to uORF translation. To examine mRNA stability in transfected cells, we sequenced mRNA reporters from cells collected at several time points after mRNA transfection. We observed a broad range of mRNA stability in transfected cells conferred by the 10-nt long 5’UTR random sequence (Fig. 2a, red line). By contrast, in cell-free lysates, these reporters showed similar stability with negligible variation (Fig. 2a, blue line). Notably, the cytoplasmic extracts maintain RNA decay activities as reported before ^16^, but not translation. The poor correlation between *in vivo* and *in vitro* half-lives of mRNA reporters supports the crucial role of translation in mRNA stability (Supplementary Fig. 4b).

**Fig. 2.**
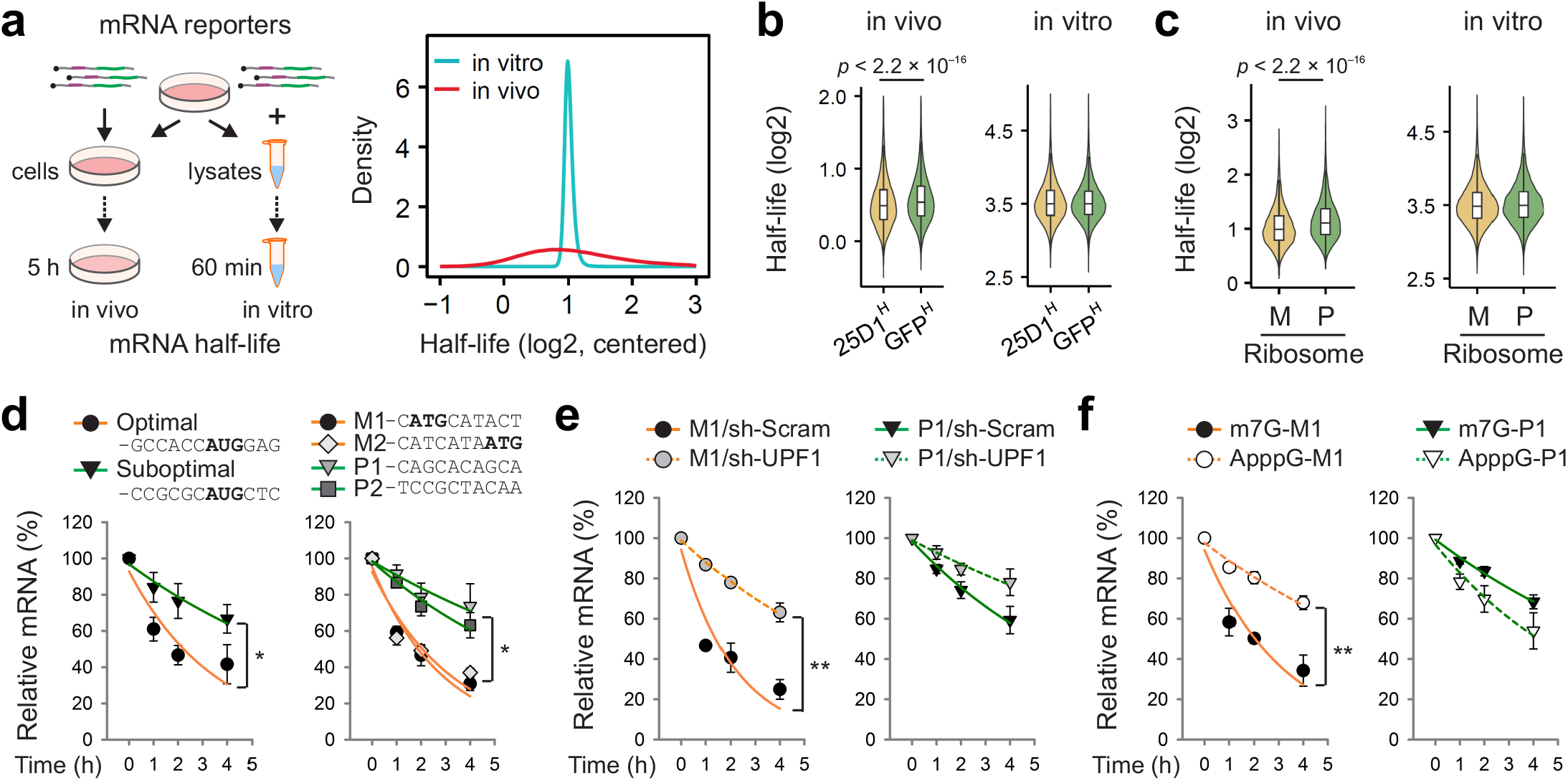
uORF translation triggers mRNA decay in a ribosome-dependent manner. **(a)** The left panel shows the schematic of measuring *in vivo* and *in vitro* half-lives of uORF reporters. The right panel shows variations of estimated *in vivo* (red) and *in vitro* (blue) half-lives (log2) with their distribution centered to the median. **(b)** Violin plots show half-lives of uORF reporters enriched in different cell populations sorted by FACS. The top 10% sequence variants ranked in 25D1^H^ and GFP^H^ populations are used for displaying *in vivo* (left) and *in vitro* (right) half-lives. Boxplots show the lower, upper quartile and the median of half-lives. P values are calculated by Wilcox test. **(c)** Violin plots show half-lives of uORF reporters enriched in different ribosome fractions. The top 10% sequences variants ranked in monosome (M) and polysome (P) are used for displaying *in vivo* (left) and *in vitro* (right) half-lives. Boxplots show the lower, upper quartile and the median of half-lives. P values are calculated by Wilcox test. **(d)** HEK293-K^b^ cells were transfected with mRNA reporters with or without optimal AUG codons (left) or representative hits from M and P fractions (right), followed by RT-qPCR at indicated time points (*n* = 3 biological replicates; *t* test). Error bars indicate SEM. * *P* < 0.05. **(e)** HEK293-K^b^ cells with or without UPF1 knockdown were transfected with representative mRNA reporters from monosome (M1, left) or polysome (P1, right) fractions, followed by RT-qPCR at indicated time points (*n* = 3 biological replicates; *t* test). Error bars indicate SEM. ** *P* < 0.01. **(f)** HEK293-K^b^ cells were transfected with representative mRNA reporters (M1, left; P1, right) with the functional m^7^G cap or non-functional ApppG, followed by RT-qPCR at indicated time points (*n* = 3 biological replicates; *t* test). Error bars indicate SEM. ** *P* < 0.01.

To relate mRNA translatability and stability, we compared the half-lives of mRNA reporters in distinct cell populations as well as different ribosome fractions. Messengers enriched in the GFP^H^ population are more stable than those in 25D1^H^ population (*p* < 2.2 × 10^−16^, Wilcox-test, Fig. 2b). The stabilizing effect of GFP translation was further validated by comparing cell populations with differential GFP intensity (Supplementary Fig. 4c). Similarly, uORF reporters enriched in polysome have significantly longer half-lives than those in monosome (*p* < 2.2 × 10^−16^, Wilcox-test, Fig. 2c). The negative role of uORF translation in mRNA stability was confirmed by individual mRNA reporters harboring optimal or suboptimal uTIS codons (Fig. 2d).

Additionally, individual mRNA decay assays of chosen uORF reporters further corroborated the notion that active uORF translation destabilizes the reporter mRNA (Fig. 2d). We observed the similar result upon plasmid transfection (Supplementary Fig. 4d).

The destabilizing effect of uORF translation is reminiscent of nonsense-mediated decay (NMD) of transcripts containing premature stop codons ^3^, although the reporters we employ do not involve the exon-junction complex (EJC) as these intron-free reporters are delivered directly to the cytosol. Since UPF1 is engaged in diverse mRNA decay pathways ^17, 18^, we measured the stability of selected uORF reporters in cells with or without UPF1 knockdown. A short-lived monosome resident (M1) with active uORF translation was significantly stabilized upon UPF1 depletion (Fig. 2e). To ensure whether the uORF-mediated mRNA decay is translation-dependent, we replaced the 5’end m^7^G cap of both M1 and P1 reporters with a non-functional cap analog ApppG. With little active translation (Supplementary Fig. 4e), the otherwise short-lived M1 was markedly stabilized by the cap analog (Fig. 2f). By contrast, the t_1/2_ of the long-lived P1 is shortened in the absence of the functional cap. These results point to opposing effects of translating ribosomes on the fate of mRNA: while GFP translation is protective, uORF translation triggers mRNA decay via a surveillance pathway involving UPF1.

The diversity of our library allowed us to identify sequences supporting neither uORF nor GFP translation by deep sequencing mRNAs present in the ribosome-free fraction. As expected, AUG was no longer enriched in this non-translatable population (Fig. 3a). Intriguingly, we observed an enrichment of GGC- and CGC-motif within the 10-nt insert (Fig. 3b). To affirm the inhibitory role of the GGC element in 5’UTR, we constructed individual mRNA reporters with each of the 8 top hits (N1 – N8). Flow cytometry confirmed low signals of both 25D1 and GFP for these variants, in contrast to M and P reporters with active translation, respectively, of uORF and GFP (Fig. 3c and Supplementary Fig. 5a). Notably, those poorly-translatable mRNA reporters exhibit significantly shorter half-lives than those engaged with ribosomes (*p* < 2.2 × 10^−16^, Wilcox-test, Fig. 3D), which was verified by individual mRNA decay assays (Supplementary Fig. 5b). These findings further strengthen the positive correlation between mRNA translatability and stability.

**Fig. 3.**
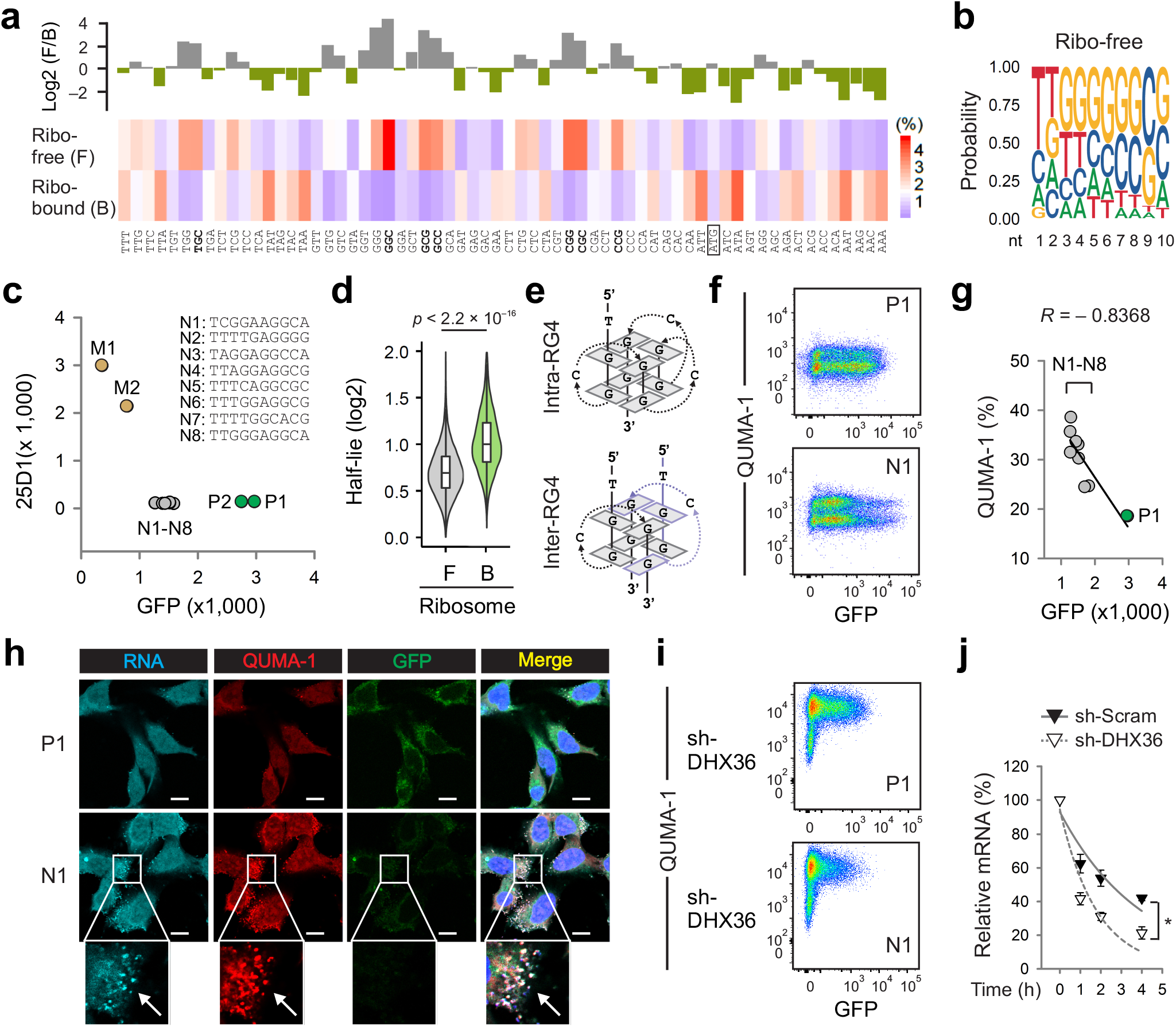
RG4 in 5’UTR triggers mRNA decay in a ribosome-independent manner. **(a)** HEK293-K^b^ cells were transfected with mRNA reporters followed by ribosome fractionation. A bar plot (top) shows the ratio of triplet frequency within the random sequences enriched in ribosome free (F) and ribosome-bound (B) fractions. The original frequency of triplets in different ribosome fractions is shown as a heat map (bottom). The GGC-like triplets are highlighted. **(b)** Sequence logo of 10 nt random sequences enriched in the ribosome free fraction. **(c)** A scatter plot shows the 25D1 and GFP fluorescence intensity of HEK293-K^b^ cells transfected with representative mRNA reporters (N1 - N8) enriched in the ribosome free fraction. **(d)** Violin plots show half-lives of uORF reporters enriched in different ribosome fractions. Boxplots show the lower, upper quartile and the median of half-lives. P values are calculated by Wilcox test. **(e)** Schematic of intramolecular (top) and intermolecular (bottom) RNA G-Quadruplexes. **(f)** Representative flow cytometry scatterplots of HEK 293-K^b^ cells transfected with P1 (top) or N1 (bottom) mRNA reporters. Transfected cells were stained with QUMA-1. **(g)** A scatter plot shows the negative correlation between the percentage of positive QUMA-1 population and GFP fluorescence intensity for mRNA reporter. **(h)** Representative confocal images of HEK 293-K^b^ cells transfected with P1 or N1 mRNA reporters. The mRNA reporters were synthesized in the presence of Alexa Fluor-UTP and the transfected cells were co-stained with QUMA-1. DNA was counter-stained with DAPI. Arrowheads indicate typical mRNA foci. Bar, 10 μm. Images are representative of at least 50 cells. **(i)** Representative flow cytometry scatterplots show the effect of DHX36 knockdown on HEK 293-K^b^ cells transfected with P1 (top) or N1 (bottom) mRNA reporters. Transfected cells were stained with QUMA-1. **(j)** HEK293-K^b^ cells with or without DHX36 knockdown were transfected with N1 mRNA reporters followed by RT-qPCR at indicated time points (*n* = 3 biological replicates; *t* test). Error bars indicate SEM. * *P* < 0.05.

We next investigated how the GGC-motif in 5’UTR impairs translation. We noticed that many sequences bearing the GGC-motif coincided with computationally predicted RNA G-quadruplex (RG4) structures ^19, 20^ (Fig. 3e). Unlike the RNA stem-loop, RG4 structures rely on non-Watson-Crick interactions between paired G quartets connected by at least one linker nucleotide (A or C). To quantitatively assess RG4 formation in live cells transfected with mRNA reporters, we used the RG4-specific fluorescent probe QUMA-1 ^21^. Unlike the P1 reporter that showed only basal levels of QUMA-1 staining, all the N reporters displayed a subpopulation with elevated QUMA-1 signals (Fig. 3f and Supplementary Fig. 5c). Intriguingly, the percentage of QUMA1 positive populations is inversely correlated to the overall GFP intensity (*R* = −0.8368, Fig. 3g). This feature implies dynamic folding and unfolding of RG4 structures *in vivo*, forming a threshold controlling RG4 translatability.

The relative instability of RG4-containing mRNA reporters is likely the result of impaired translation. Unlike the short-lived M1 reporter, the t_1/2_ of the non-translatable N1 is insensitive to UPF1 knockdown (Supplementary Fig. 6a). We systematically examined the role of other decay factors and found that N1 is mostly stabilized in cells lacking either DCP2 or XRN1 (Supplementary Fig. S6b), but not CNOT1 or PAN3 (Supplementary Fig. S6c). This result suggests that the degradation of RG4 mRNA follows the 5’ → 3’ decay pathway. To probe the degradation pathway in more detail, we tracked the intracellular localization of RG4 reporters in live cells by synthesizing individual mRNA reporters using Alexa Fluo-labeled UTP. Compared to the P1 reporter that distributed uniformly in the cytoplasm, the RG4-containing N1 reporters form distinct foci co-stained with QUMA-1 (Fig. 3h). Additionally, these foci overlaps with the P-body marker DCP2 (Supplementary Fig. 6d), further supporting a distinct decay pathway for RG4 mRNAs.

RG4 also exists in endogenous transcripts as exemplified by the oncogene *NRAS* ^22^. We confirmed that the wild type 5’UTR of *NRAS* shortened the t_1/2_ of the uORF reporter when compared to the mutant lacking RG4 (Supplementary Fig. 7a). Several helicases such as DHX36 are known to unwind RG4 structures, permitting mRNA translation ^23^. Consistently, knocking down DHX36 resulted in marked accumulation of QUMA-1 signals with further reduced GFP levels in transfected cells (Fig. 3i). Importantly, DHX36 depletion accelerates the decay of N1 reporters (Fig. 3j) as well as the endogenous *NRAS* (Supplementary Fig. 7b). Collectively, the RG4 in 5’UTR not only hampers ribosome scanning, but also actively targets mRNA for degradation in the P-body.

We next explored whether certain sequence variants in 5’UTR enable cap-independent translation. Unlike previous efforts using bi-cistronic constructs ^24^, we synthesized an entire mRNA library capped with ApppG, which does not support cap-dependent translation in transfected cells (Supplementary Fig. 4e). We separated the ribosome-associated fraction from the ribosome free fraction followed by deep sequencing. Despite the lack of a functional cap, a substantial amount of variants were recovered from the ribosome fractions (Fig. 4a). Remarkably, the ribosome-bound mRNA reporters are enriched with an A-rich sequence element in 5’UTR (Fig. 4b). By contrast, a C-rich element is overrepresented in the ribosome-free fraction. As an independent validation, we constructed individual mRNA reporters with the insert containing a string of 10A or 10C. In the presence of the functional m^7^G cap, both 10A and 10C exhibit strong GFP signals (Fig. 4c). In the presence of the ApppG cap analog, however, only 10A shows robust GFP translation. Since GFP is located about 80 nt downstream of the insert, the 10A-mediated cap-independent translation is compatible with ribosome scanning.

**Fig. 4.**
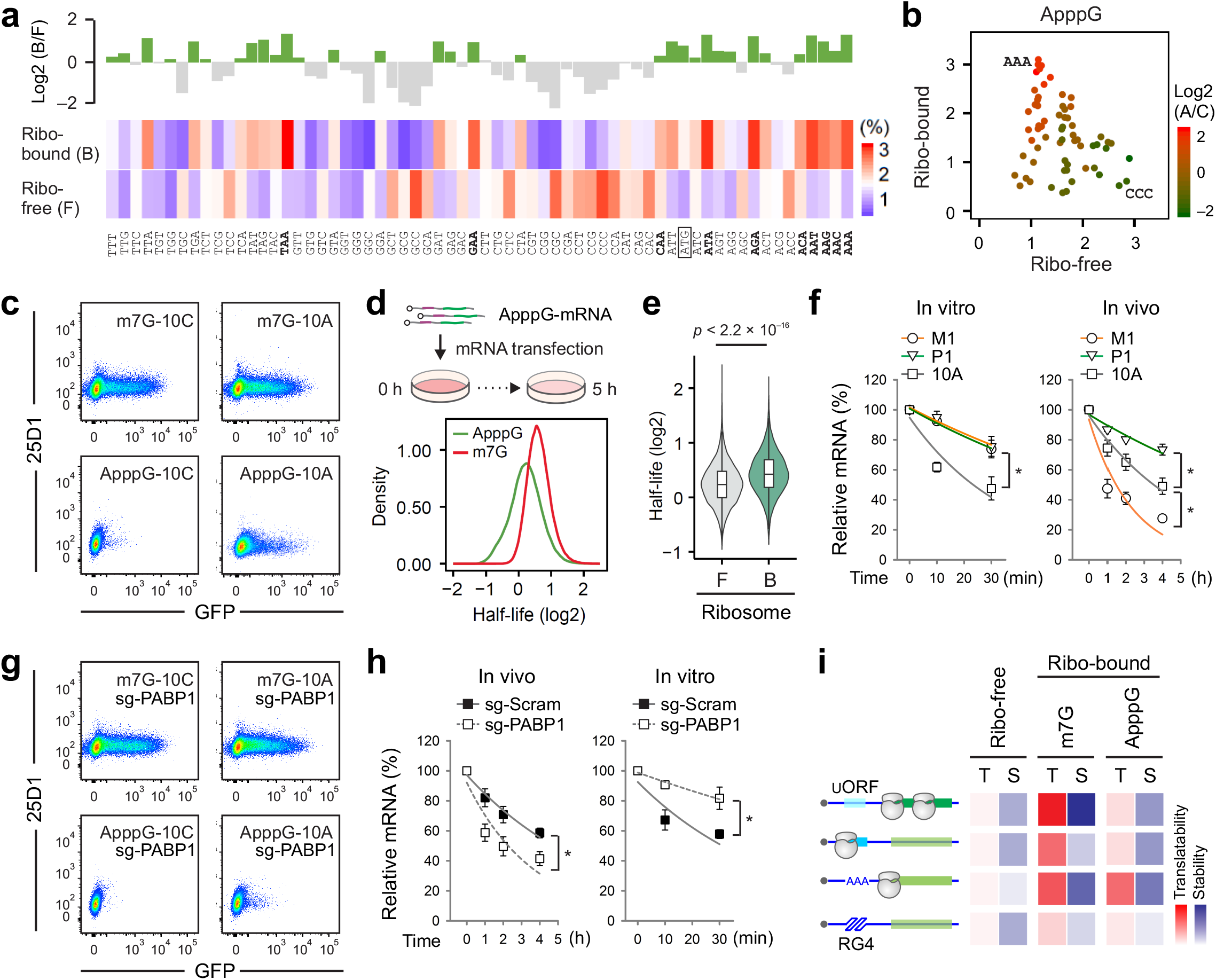
An A-rich element in 5’UTR promotes cap-independent translation and translation-independent decay. **(a)** HEK293-K^b^ cells were transfected with mRNA reporters capped with ApppG followed by ribosome fractionation. A bar plot (top) shows the ratio of triplet frequency within the random sequences enriched in ribosome free (F) and ribosome-bound (B) fractions. The original frequency of triplets in different ribosome fractions is shown as a heat map (bottom). The A-rich triplets are highlighted. **(b)** A scatter plot shows the distribution of triplets between ribosome free and ribosome-bound fractions. Note the clear segregation between A-rich and C-rich triplets. All points are color-encoded based on the ratio of A/C. **(c)** Representative flow cytometry scatterplots of HEK 293-K^b^ cells transfected with 10C (left) or 10A (right) mRNA reporters capped with m^7^G (top) or ApppG (bottom). **(d)** The top panel shows the schematic of measuring *in vivo* half-lives of mRNA reporters capped with ApppG. The bottom panel shows variations of estimated half-lives (log2) for mRNA reporters capped with m^7^G (red) or ApppG (green). **(e)** Violin plots show half-lives of uORF reporters capped with ApppG enriched in different ribosome fractions. Boxplots show the lower, upper quartile and the median of half-lives. P values are calculated by Wilcox test. **(f)** The decay of mRNA reporters in the lysates of HEK293-K^b^ cells (left) was determined by RT-qPCR at indicated time points. For *in vivo* decay, HEK293-K^b^ cells were transfected with mRNA reporters followed by RT-qPCR at indicated time points (*n* = 3 biological replicates; *t* test). Error bars indicate SEM. * *P* < 0.05. **(g)** Representative flow cytometry scatterplots show the effect of PABP1 knockdown on HEK 293 cells transfected with 10C (left) or 10A (right) mRNA reporters capped with m^7^G (top) or ApppG (bottom). **(h)** HEK293 cells with or without PABP1 knockdown were transfected with 10A mRNA reporters followed by RT-qPCR at indicated time points (left). The decay of 10A mRNA reporters in the lysates of HEK293-K^b^ cells with or without PABP1 knockdown (right) was determined by RT-qPCR at indicated time points (*n* = 3 biological replicates; *t* test). Error bars indicate SEM. * *P* < 0.05. **(i)** A summary of diverse mechanisms linking translation initiation and mRNA stability. mRNA translatability is color-coded as red, whereas stability as blue. mRNAs are stratified based on uORF translation or 5’UTR sequence elements. Different translation modes (ribosome-free, cap-dependent, and cap-independent) are also considered.

The mRNA library capable of cap-independent translation offers another means of assessing the relationship between translatability and stability. We systematically measured the stability of ApppG-capped mRNA reporters in transfected cells (Fig. 4d). Not surprisingly, the non-functional cap analog reduces the overall half-life of messages in comparison to the m^7^G-capped counterparts. Further supporting the stabilizing effect of translating ribosomes, the sequence variants capable of cap-independent translation are significantly more stable than the non-translatable ones (*p* < 2.2 × 10^−16^, Wilcox-test, Fig. 4e). Indeed, in the presence of the cap analog ApppG, 10A becomes more stable than 10C in transfected cells (Supplementary Fig. 8a). By contrast, the stability of both reporters bearing the functional m^7^G cap is comparable.

We noticed that the A-rich element appears to be underrepresented in transfected cells with positive 25D1 or GFP signals (Supplementary Fig. 2d). One possibility is the faster turnover of these mRNAs before ribosome engagement. An inspection of *in vitro* mRNA stability uncovered an enrichment of the poly(A) tract from mRNA reporters short-lived in cell lysates (Supplementary Fig. 8b). Consistently, 10A exhibits a much faster turnover rate than M1 and P1 in the absence of translation (Fig. 4f). The *in vitro* destabilizing effect of A-rich sequence is further supported by variants bearing different amount of A residues (Supplementary Fig. 8c). To investigate the *in vitro* decay mechanisms conferred by the poly(A) tract in 5’UTR, we prepared lysates from cells lacking individual decay factors. Depletion of UPF1 showed little effect on the turnover of 10A *in vivo* or *in vitro* (Supplementary Fig. 8d). However, lacking CNOT1, PARN, or PAN3 stabilized 10A, but not M1 and P1, in the lysates (Supplementary Fig. 8e). Both Ccr4-Not and Pan2-Pan3 complexes participate in the 3’ end poly(A) tail shortening ^25^. It is likely that the poly(A) tract in 5’UTR follows the similar mechanism to degrade mRNAs.

An unstructured A-rich element in 5’UTR has been shown to promote cap-independent mRNA translation via PABP1 ^26^. Indeed, knocking down PABP1 eliminates the GFP signals from the 10A reporter capped with ApppG (Fig. 4g). Notably, PABP1 depletion had little effects on the translation of mRNAs capped with m^7^G. Intriguingly, PABP1 depletion destabilized the 10A reporter in transfected cells but stabilized the same reporter in the cell lysates (Fig. 4h). These results suggest a dichotomy in mRNA surveillance pathways: while the interaction between poly(A) and PABP1 protects mRNA from degradation by enabling cap-independent translation, it promotes mRNA decay when translation is inactive.

## Discussion

Our results indicate that the multi-faceted mRNA surveillance system starts from 5’ leader even before the ribosome engagement. Although the role of 3’UTR in mRNA decay has long been appreciated ^27^, recent studies suggest that ribosome-mediated mRNA surveillance pathways primarily act on CDS ^3, 4^. The potential role of 5’UTR in mRNA stability remained elusive until now. The RNA-based uORF reporter system we devised here establishes the functional correlation between mRNA translatability and stability in an unbiased manner. With identical coding sequence and 3’UTR, it is remarkable that a 10 nt sequence variation in 5’UTR controls mRNA translatability and stability by an order of magnitude. In addition to revealing the sequence principles in forming TIS sites, we uncovered a subset of sequence variants that block ribosome scanning. We further probed 5’UTR sequence elements that enable cap-independent translation. Importantly, the poly(A) tract in 5’UTR acts like the internal ribosome entry site (IRES) but permits the scanning process for the recognition of downstream start codons.

Perhaps our most surprising finding is the diverse mechanistic connection between translation initiation and mRNA decay (Fig. 4i). Although translating ribosomes protects mRNA from degradation in general, uORF translation triggers mRNA decay via UPF1. RG4 in 5’UTR impairs ribosome scanning and relocates the messenger to the P-body, forming an example of ribosome-independent mRNA decay mechanism. Since not all the messengers are simultaneously engaged with ribosomes inside cells, a given mRNA molecule could experience differential stability. Indeed, the poly(A) leader enables cap-independent mRNA translation by recruiting PABP1, thereby enhancing its stability. However, it also destabilizes the same mRNA when translation is inactive. The 5’ leader-initiated mRNA surveillance pathways suggest a broad quality control network monitoring mRNA structures, interacting factors, and ribosome engagement. Deciphering the regulatory code embedded in 5’UTR will enable more accurate prediction of translational output and promises improved design of sequence elements to optimize protein expression in synthetic biology.

## Acknowledgements

We thank the Grimson lab for providing us several shRNAs targeting mammalian decay factors. We are grateful to Cornell University Life Sciences Core Laboratory Center for sequencing, FACS, and confocal microscope support. This work was supported by US National Institutes of Health (R01GM1222814 and R21CA227917) and HHMI Faculty Scholar (55108556) to S.-B.Q.

## Author Contributions

S.-B.Q. conceived the project and designed the experiments. L.J. performed the majority of experiments and Y.M. conducted the majority of data analysis. Q.J. contributed to the PABP1 knockdown experiments. D.D. and J.W.Y. helped with 25D1 reagents and HEK293-K^b^ cells. S.-B.Q. wrote the manuscript with comments from L.J. and Y.M. All authors discussed the results and edited the manuscript.

## Supplementary Information

### Supplementary Figures

**Fig. 1.**
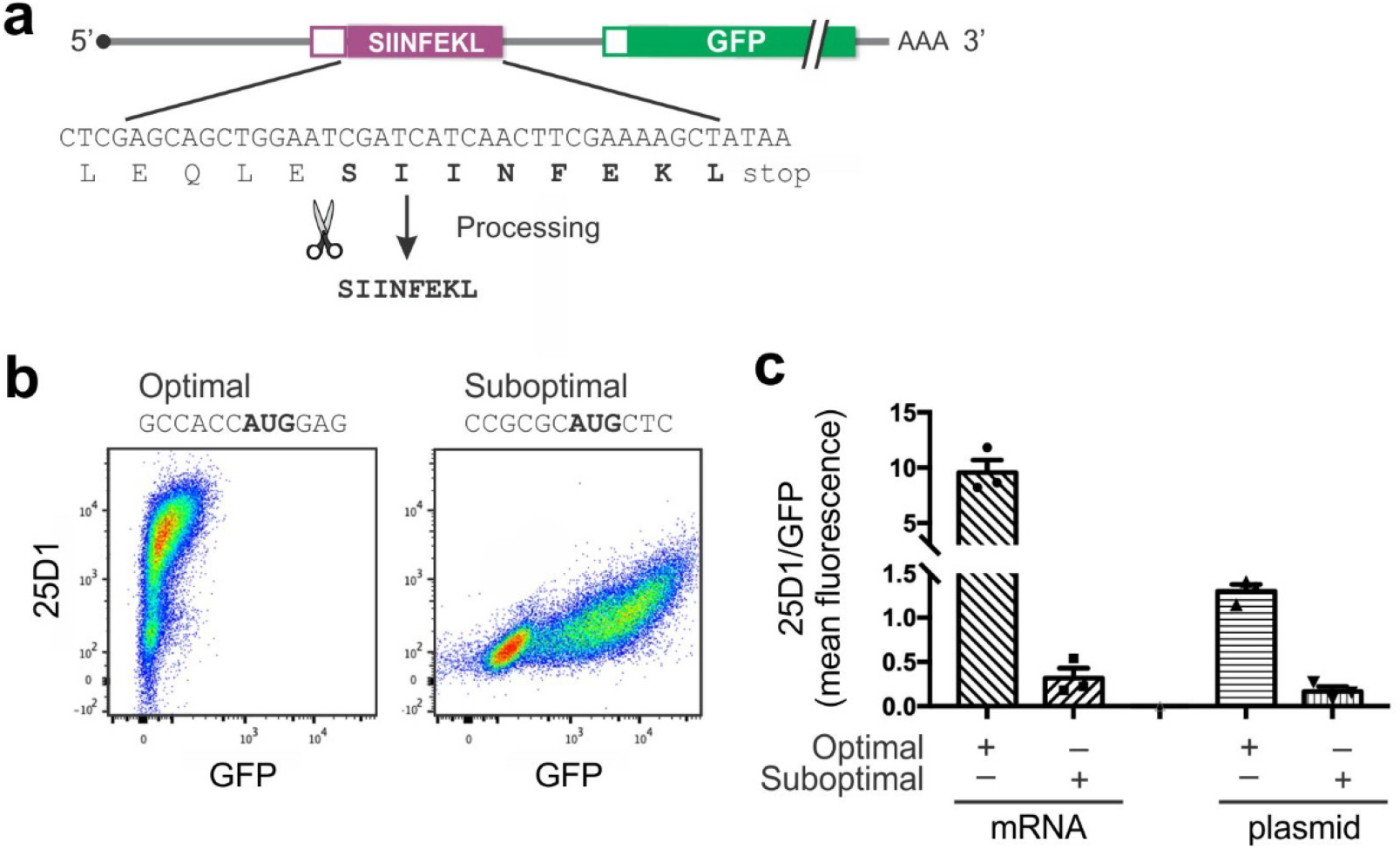
Characterization of uORF reporters. **(a)** Basic design of the uORF reporter with the SIINFEKL sequence highlighted. 5 additional amino acids (LEQLE) are present, which permits processing of SIINFEKL from the same flanking amino acids regardless of the TIS sequence. **(b)** Representative flow cytometry scatterplots of HEK 293-K^b^ cells transfected with synthetic mRNA reporters with optimal or suboptimal AUG codons. **(c)** A bar graph shows the ratio of 25D1/GFP in HEK 293-K^b^ cells transfected with synthetic mRNA or plasmid DNA. Error bars, mean ± s.e.m; *n* = 3 biological replicates.

**Fig. 2.**
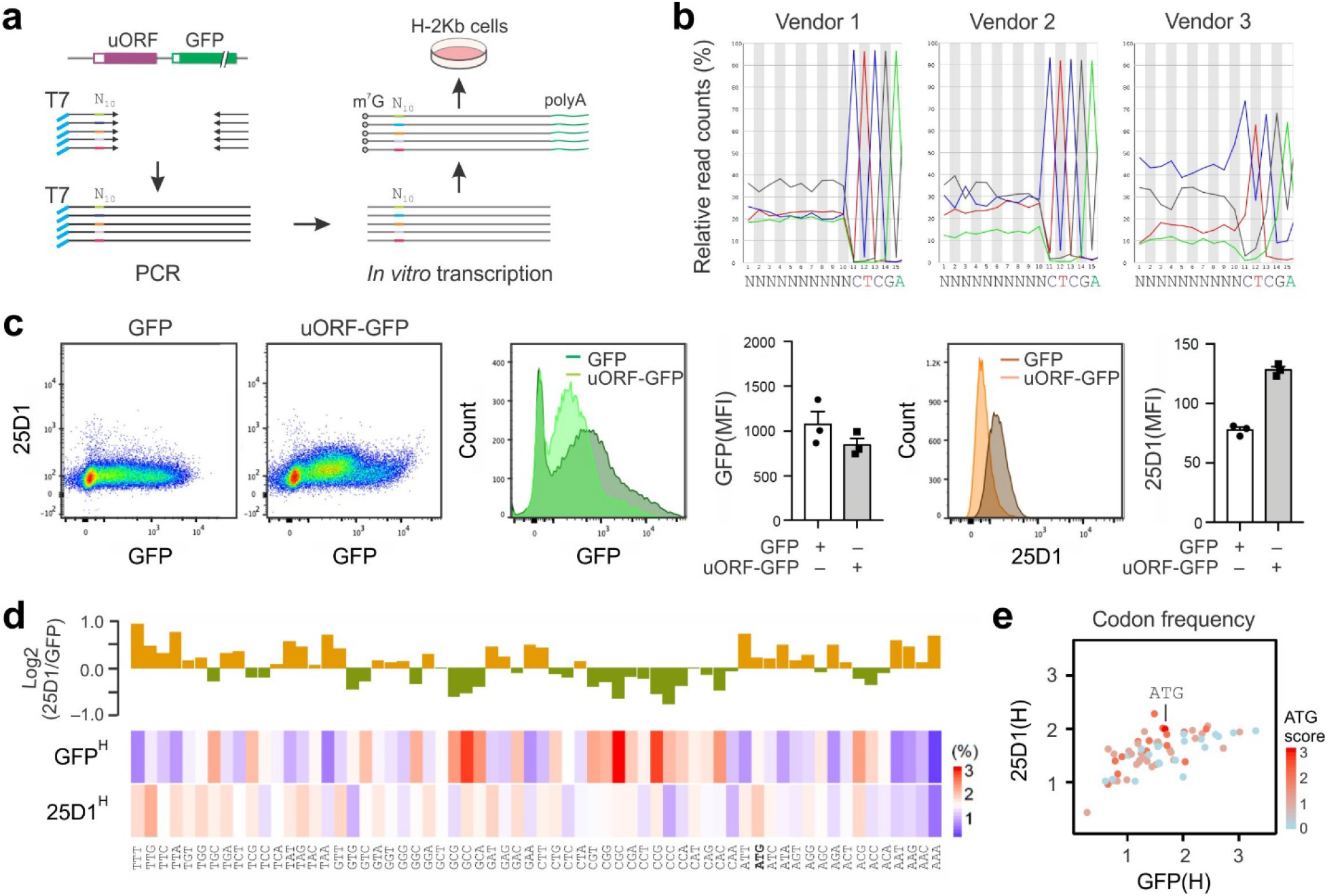
Establishing massively parallel uORF reporters. **(a)** Schematic of generating a library of RNA-based uORF reporters by PCR-amplification using primers composed of random 10-nt sequences upstream of the uORF. Pooled PCR products were utilized as templates for in vitro RNA synthesis followed by 5’ capping and 3’ polyadenylation. **(b)** Comparison of sequence randomness for nucleotide oligos synthesized by different vendors. **(c)** Representative flow cytometry scatterplots of HEK 293-K^b^ cells transfected with the GFP mRNA reporter or pooled uORF reporters. Relative GFP and 25D1 fluorescence intensity between GFP and uORF-GFP reporters are shown in histograms as well as bar graphs. Error bars, mean ± s.e.m. n = 3 biological replicates. **(d)** HEK293-K^b^ cells were transfected with mRNA reporters followed by FACS soring into GFP^H^ and 25D1^H^ populations. A bar plot (top) shows the ratio of triplet frequency within the random sequences enriched in the 25D1^H^ population over the GFP^H^ population. Only the top 10% sequence variants ranked in 25D1^H^ and GFP^H^ populations are used. The original frequency of triplets in different populations is shown as a heat map (bottom). **(e)** Correlation of triplet frequencies within the sequence variants enriched in 25D1^H^ or GFP^H^ populations. All points are color-encoded based on the similarity to ATG.

**Fig. 3.**
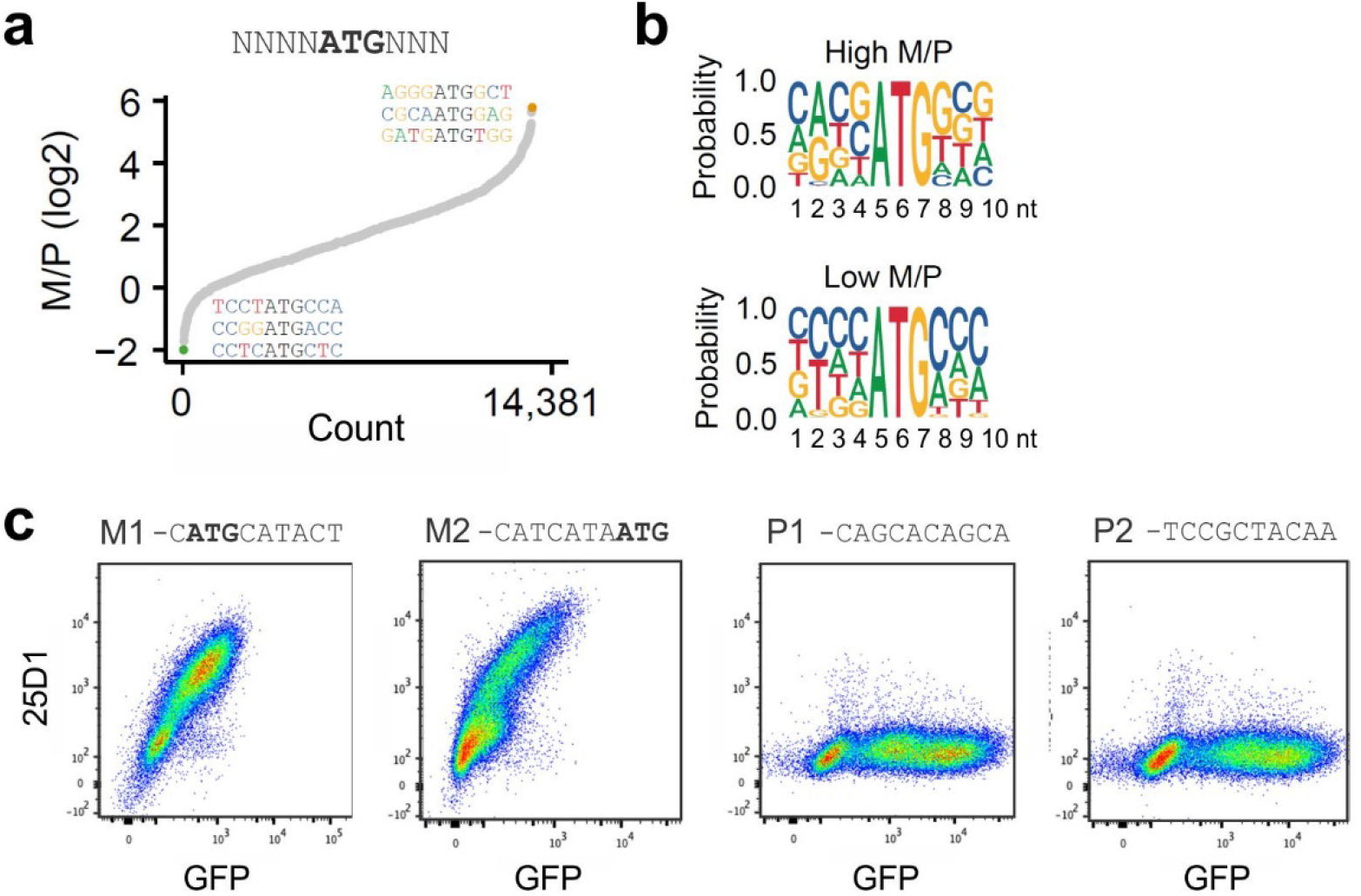
Identification of TIS sequence features in uORF reporters enriched in ribosome fractions. **(a)** A total of 14,381 sequences with all permutations of NNNNAUGNNN are ranked based on the ratio of frequency between monosome and polysome. Both the top and bottom hits are highlighted. **(b)** Sequence logo of 10 nt random sequences with high (top) or low (bottom) M/P ratio. Note that the high M/P sequence is consistent with the Kozak consensus sequence. **(c)** Representative flow cytometry scatterplots of HEK 293-K^b^ cells transfected with mRNA reporters with sequence variants chosen from monosome (M1, M2) or polysome (P1, P2).

**Fig. 4.**
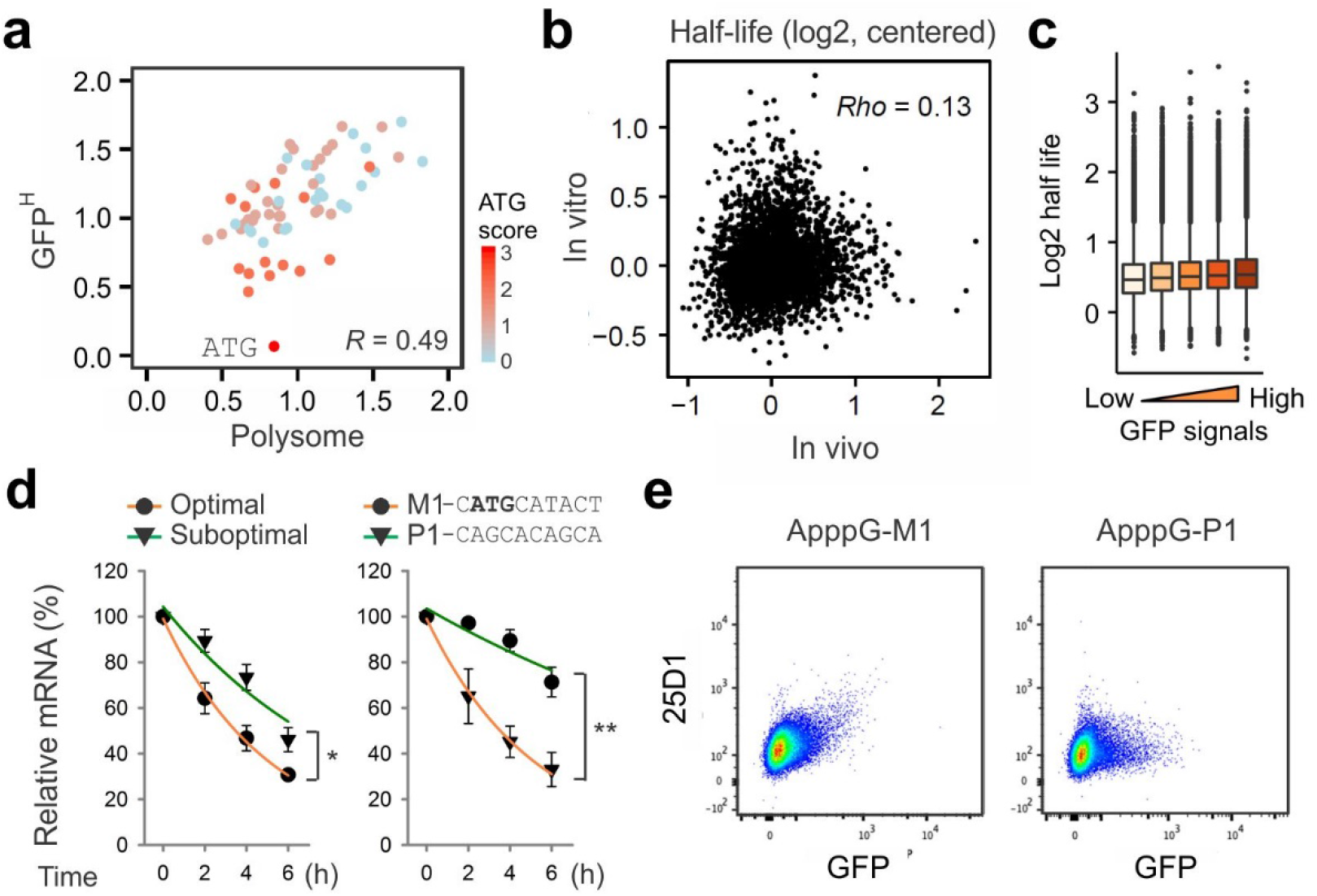
uORF translation triggers mRNA decay in a ribosome-dependent manner. **(a)** A scatter shows the correlation of triplet frequencies enriched in polysome and GFP^H^ population from HEK293-K^b^ cells transfected with mRNA reporters. All points are color-encoded based on the similarity to ATG. **(b)** A scatter shows the correlation of *in vivo* and *in vitro* half-lives of mRNA reporters. Half-life values were centered to medians. **(c)** A boxplot shows positive correlation between GFP intensities and half-lives of mRNA reporters. All random sequences were divided into five groups based GFP intensity measured by flow cytometry. **(d)** HEK293-K^b^ cells were transfected with DNA plasmids with or without optimal ATG codons (left) or representative hits from M and P fractions (right), followed by RT-qPCR at indicated time points (*n* = 3 biological replicates; *t* test). Error bars indicate SEM. ** *P* < 0.01; * *P* < 0.05. **(e)** Representative flow cytometry scatterplots of HEK 293-K^b^ cells transfected with mRNA reporters capped with ApppG with sequence variants chosen from monosome (M1) or polysome (P1).

**Fig. 5.**
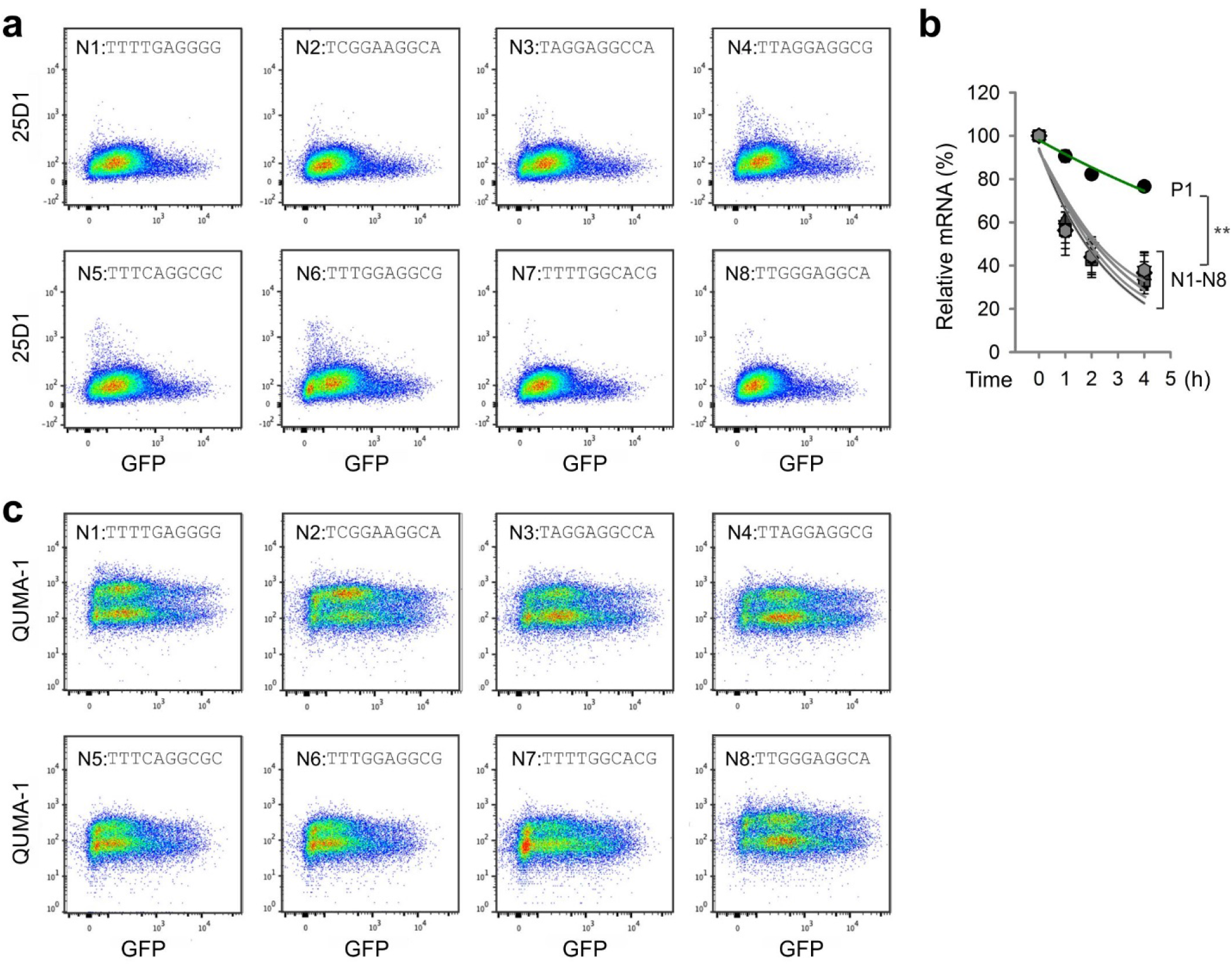
Identification of inhibitory elements in 5’UTR that impair translation. **(a)** Flow cytometry scatterplots of HEK 293-K^b^ cells transfected with mRNA reporters enriched in the ribosome-free fractions (N1 – N8). **(b)** HEK293-K^b^ cells were transfected with mRNA reporters with sequence variants chosen from the ribosome-free fractions (N1 – N8), followed by RT-qPCR at indicated time points (*n* = 3 biological replicates; *t* test). Error bars indicate SEM. ** *P* < 0.01. **(c)** Flow cytometry scatterplots of HEK 293-K^b^ cells transfected with mRNA reporters enriched in the ribosome-free fractions (N1 - N8) and stained with QUMA-1.

**Fig. 6.**
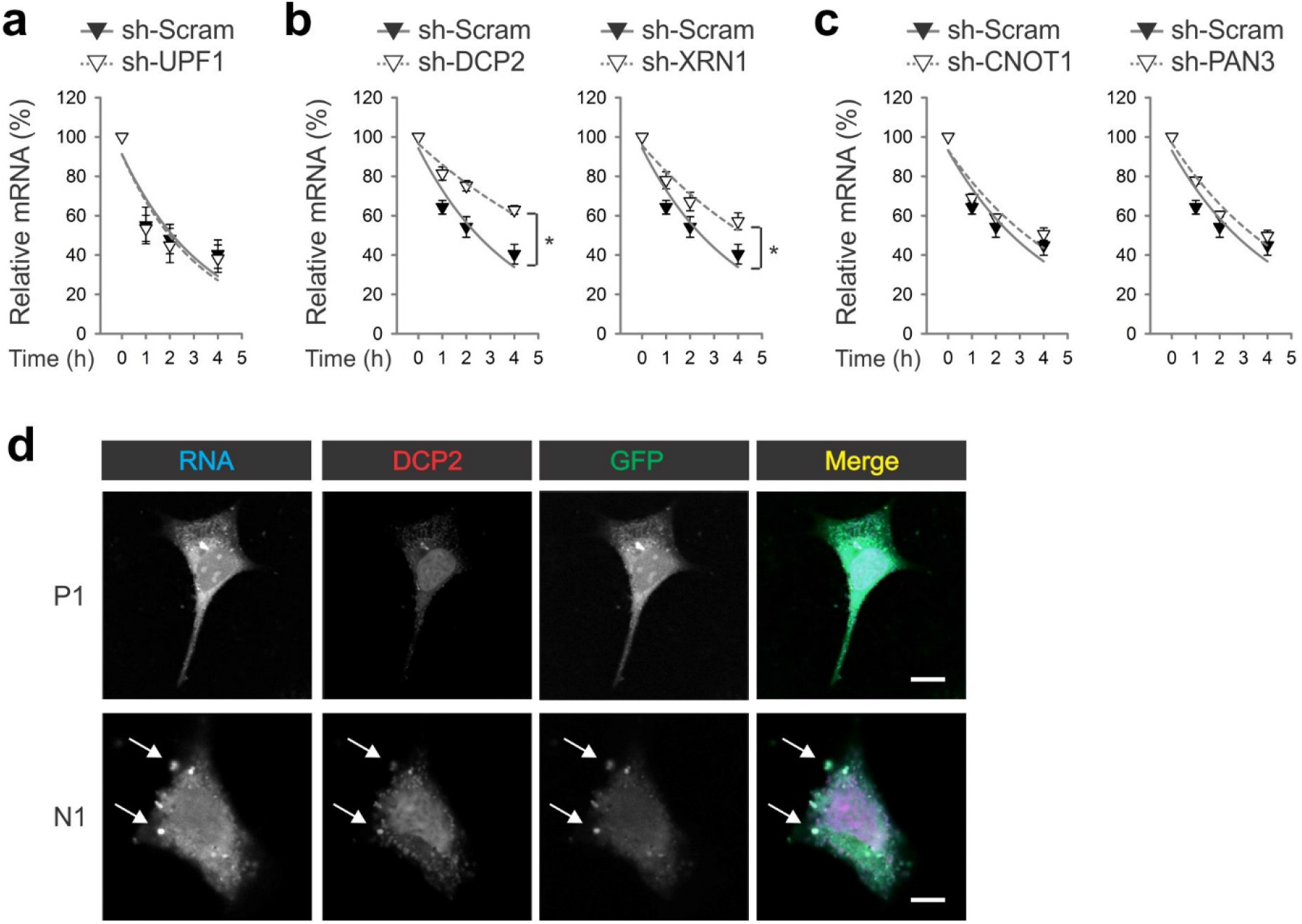
RG4 in 5’UTR triggers mRNA decay in the P-body. **(a)** HEK293-K^b^ cells with or without UPF1 knockdown were transfected with the N1 mRNA reporter, followed by RT-qPCR at indicated time points. (*n* = 3 biological replicates; *t* test). Error bars indicate SEM. **(b)** HEK293-K^b^ cells with DCP2 (left) or XRN1 (right) knockdown were transfected with the N1 mRNA reporter, followed by RT-qPCR at indicated time points. (*n* = 3 biological replicates; *t* test). Error bars indicate SEM. * *P* < 0.05. **(c)** HEK293-K^b^ cells with CNOT1 (left) or PAN3 (right) knockdown were transfected with the N1 mRNA reporter, followed by RT-qPCR at indicated time points. (*n* = 3 biological replicates; *t* test). Error bars indicate SEM. **(d)** Representative confocal images of HEK 293-K^b^ cells transfected with P1 or N1 mRNA reporters. The mRNA reporters were synthesized in the presence of Alexa Fluor-UTP and the transfected cells were co-stained with a DCP2 antibody. DNA was counter-stained with DAPI. Arrowheads indicate typical mRNA foci. Bar, 10 μm. Images are representative of at least 50 cells.

**Fig. 7.**
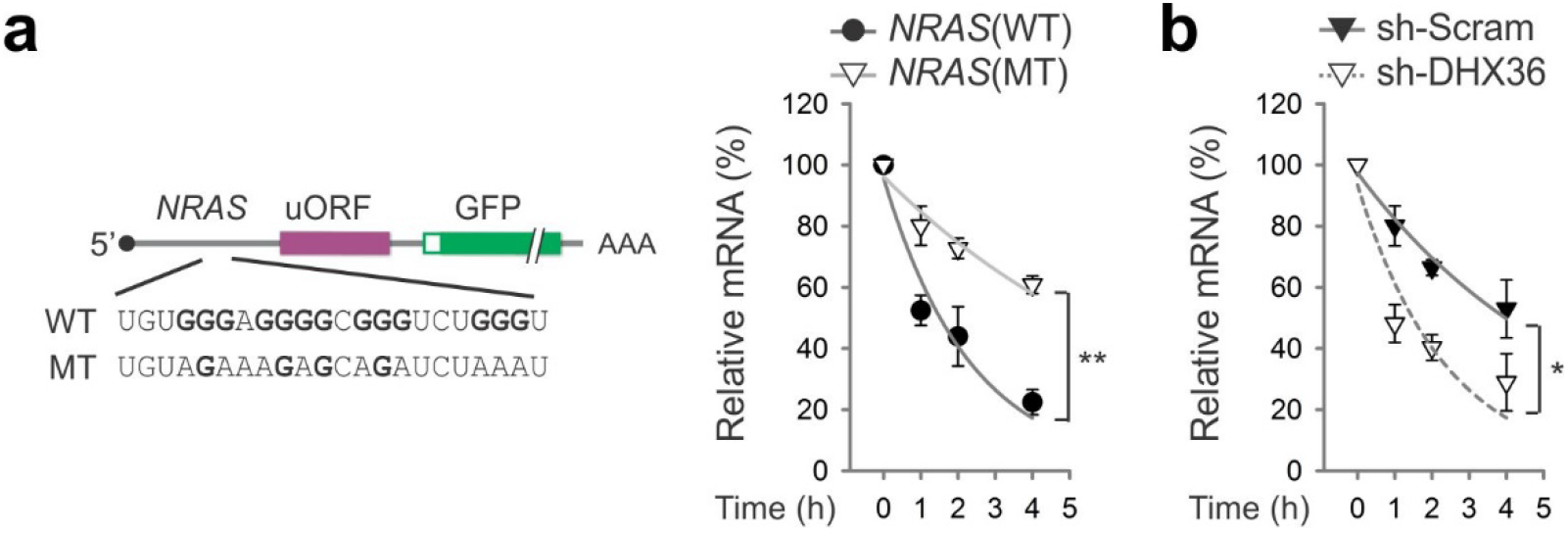
RG4 in 5’UTR derived from *NRAS* triggers mRNA decay. **(a)** The left panel shows the schematic of mRNA reporter with 5’UTR derived from *NRAS* with (WT) or without (MT) RG4. The right panel shows the decay of mRNA reporters in transfected HEK293-K^b^ cells. (*n* = 3 biological replicates; *t* test). Error bars indicate SEM. ** *P* < 0.01. **(b)** The stability of endogenous *NRAS* was measured in HEK293-K^b^ cells with or without DHX36 knockdown. (*n* = 3 biological replicates; *t* test). Error bars indicate SEM. * *P* < 0.05.

**Fig. 8.**
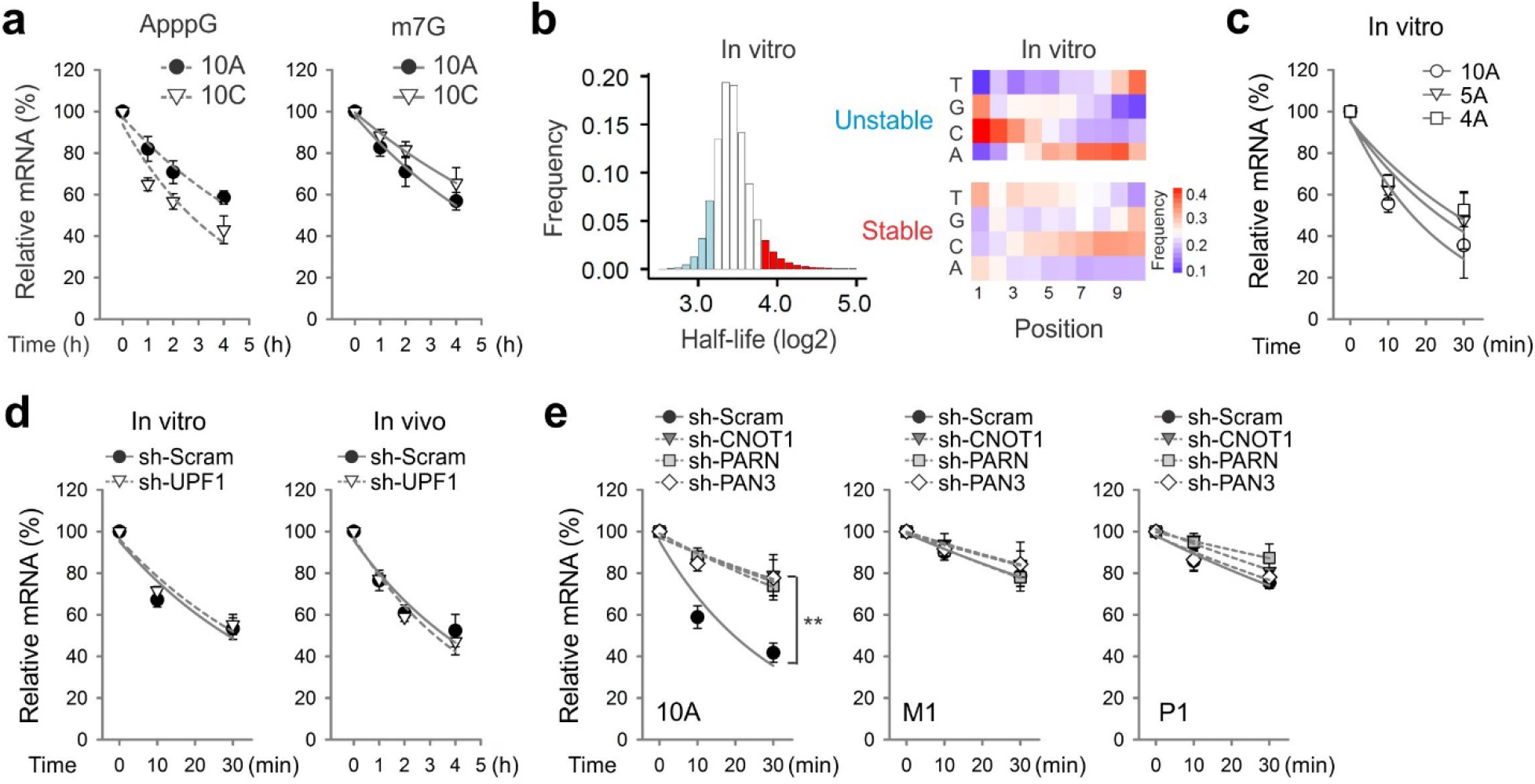
An A-rich element in 5’UTR promotes translation-independent decay. **(a)** HEK293-K^b^ cells were transfected with 10A or 10C mRNA reporters capped with m^7^G (right) or ApppG (left), followed by RT-qPCR at indicated time points. (*n* = 3 biological replicates; *t* test). Error bars indicate SEM. **(b)** The left panel shows the distribution of *in vitro* half-lives of mRNA reporters. The most stable (top 10%) sequences are highlighted in red, and the most unstable sequences are highlighted in light blue. The right panel shows the heat map of base frequency at different positions of random sequences. **(c)** The *in vitro* decay of mRNA reporters (10A, 5A, and 4A) in the lysates of HEK293-K^b^ cells was determined by RT-qPCR at indicated time points. (*n* = 3 biological replicates; *t* test). Error bars indicate SEM. **(d)** The *in vitro* decay of 10A mRNA reporters in the lysates of HEK293-K^b^ cells with or without UPF1 knockdown was determined by RT-qPCR at indicated time points (left). For the *in vivo* stability, HEK293-K^b^ cells with or without UPF1 knockdown were transfected with 10A mRNA reporters followed by RT-qPCR at indicated time points (right). (*n* = 3 biological replicates; *t* test). Error bars indicate SEM. **(e)** The *in vitro* stability of mRNA reporters (10A, M1, and P1) in the lysates of HEK293-K^b^ cells with CNOT1, PARN, or PAN3 knockdown was determined by RT-qPCR at indicated time points. (*n* = 3 biological replicates; *t* test). Error bars indicate SEM. ** *P* < 0.01.

### Methods

#### Cell lines and reagents

HEK293 cells and HEK293-K^b^ cells are cultured in Dulbecco’s Modification of Eagle’s Medium (Corning 10-013-CV) supplemented with 10% fetal bovine serum (Sigma 12306C). Anti-DCP2 (Abcam ab28658) antibody is used in the immunofluorescence staining. QUMA-1 (Millipore, SCT056) is used in RNA G-Quadruplexes staining.

#### Plasmids construction

A two-step PCR amplification approach is used to generate uORF reporters. First, full length EGFP is amplified from pcDNA3-EGFP using forward primer oligo 1 and reverse primer oligo 2 (Table S1), generating LEQLE-SIINFEKL-GFP. The resulting PCR product is used as a template to produce the full length reporter using forward primer oligo 3-34 containing 5’UTR of β-globin and reverse primer oligo 2. PCR products are cloned into pcDNA3.1 (Invitrogen) using HindIII and PmeI restriction sites to generate plasmids with optimal or suboptimal sequence context, AUG or non-AUG codons, as well as other chosen sequences.

#### In vitro transcription

To generate mRNAs suitable for transfection, 3 μg PCR products described above are utilized for in vitro transcription using T7 mMessage mMachine kit (Ambion) followed by poly(A) tailing kit (Ambion) according to the manufacturer’s instructions. mRNAs with non-functional cap analogue GpppA (NEB) are synthesized using MEGAscript T7 Transcription Kit (Ambion). Fluorescently-labeled mRNAs are synthesized using MEGAscript T7 Transcription Kit in the presence of ChromaTide Alexa Fluo 546-14-UTP (Invitrogen) according to the manufacturer’s instructions. Synthesized mRNAs are purified by Quick Spin RNA Columns (Roche).

#### Transfection

For a 6-well plate, 3 μg mRNA per well in 200 μl Opti-MEM (Life Technologies) is mixed with Lipofectamine MessengerMAX (Invitrogen) in 200 μl Opti-MEM at room temperature for 10-20 min. For a 10-cm dish, a total of 10 μg mRNA is used with a 1:1 ratio of RNA to Lipofectamine. The mixture is added to cells and incubated at 37°C for 5 h. For plasmid transfection, 2 μg of DNA per well is mixed with Lipofectamine 2000 with a 1:2 ratio of DNA (μg) to Lipofectamine 2000 (μl). The mixture is added to cells and incubated at 37°C for for 24 h.

#### Flow cytometry

Transfected HEK293-K^b^ cells are washed with PBS and harvested by trypsin. Cells are then re-suspended in blocking buffer (1% bovine serum albumin (BSA) in PBS). Cells are aliquoted into a 96-well plate followed by 2000 rpm spinning for 2 min. After removal of blocking buffer, cells are washed one more time followed by staining with 25D1 Alexa 647 antibody (1:1000 in 75 uL solution per well). After incubation in the dark with gentle rocking at 4°C for 30 minutes, cells are washed three times with 200 uL of the blocking buffer to remove unbound antibodies. Resuspend cells in 300 uL of blocking buffer followed by single cell filtering (Falcon). Cells are analyzed on a BD FACSAria Fusion flow cytometer (BD Biosciences). Cytometry data analysis is conducted using FlowJo.

#### Cell sorting

HEK293-K^b^ cells transfected with the mRNA reporter library are sorted based on the fluorescence intensity of 25D1 (Alexa Fluor 647) and GFP using an BD FACSAria Fusion flow cytometer. Cells are sorted into 4 gates: low 25D1 signal (25D1^L^), high 25D1 signal (25D1^H^), low GFP signal (GFP^L^), high GFP signal (GFP^H^). For a negative control, cells are transfected with Lipofectamine MessengerMAX only. Total RNAs from sorted cells are extracted using Trizol reagent (Invitrogen). Purified RNAs are used for cDNA library construction and high-throughput sequencing described below.

#### Polysome profiling

Sucrose solutions are prepared in polysome buffer (10 mM HEPES, pH 7.4, 100 mM KCl, 5 mM MgCl_2_ and 100 μg/ml cycloheximide). Sucrose density gradients (15-45 % (wt/vol) is freshly prepared in a SW41 ultracentrifuge tube (Backman) using a Gradient Master (BioComp Instruments). 10 μg of RNA reporter library per 10-cm dish is used for transfection for 5 h using Lipofectamine MessengerMAX. Transfected cells are then washed and lysed in polysome lysis buffer (polysome buffer with 2 % Triton X-100). Cell debris are removed by centrifugation at 14,000 rpm for 10 min at 4 °C. 500 μL of supernatant is loaded onto sucrose gradients followed by centrifugation for 2 h 30 min at 38,000 rpm 4 °C in a SW41 rotor. Separated samples are fractionated at 0.75 ml/min through an automated fractionation system (Isco) that continually monitors OD254 values. An aliquot of ribosome fractions representing ribosome-free, monosome, or polysome are collected followed by extraction of total RNAs using Trizol LS reagent (Invitrogen). Purified RNAs are used for cDNA library construction and high-throughput sequencing described below.

#### mRNA stability analysis in vivo

HEK293-K^b^ cells transfected with mRNA reporter library are collected at various time points (2, 5 h) following 1 h of transfection. Total RNAs are extracted using Trizol reagent. Purified RNA samples are used for cDNA library construction and high-throughput sequencing described below. For individual mRNA reporter half-life assay, transfected cells are collected at time points (0, 1, 2 and 4 h) followed by total RNA extraction using Trizol reagent. For half-life assay using plasmid reporters, 24 h transfected cells are treated with transcription inhibitor actinomycin D (5 μg/ml, Sigma-Aldrich) and collected at different times (0, 2, 4 and 6 h). For half-life assay of endogenous mRNAs (such as *NRAS*), the cells are treated with 5 μg/ml actinomycin D for various times (0, 1, 2 and 4 h) followed by total RNA extraction. Firefly luciferase mRNA synthesized in vitro is mixed with the total RNA samples as spike-in controls.

#### mRNA stability analysis in vitro

Cytoplasmic extracts are prepared from HEK293-K^b^ cells as described previously (*1*). In brief, cells with 80% confluence are harvested into ice-cold PBS by scrapping. Cells are re-suspended in 200 μL of cold buffer A (10 mM Hepes (pH 7.9), 1.5 mM MgCl_2_, 10 mM KCl, 1 mM DTT, and 1 mM PMSF). Pellet the cells at 4500g for 2 min at 4 °C and remove the supernatant. Cells are re-suspended in 120 μL of buffer A and transfer the suspension into a 1-mL Dounce homogenizer. Lyse the cells with 10 strokes of a B-type pestle. Transfer the lysed cells to a new tube and pellet the nuclei at 4500 g for 2 min at 4 °C. Transfer the supernatant to a new tube and mix the cytoplasmic fraction with 0.11 vol of ice-cold buffer B (0.3 M Hepes (pH 7.9), 1.4 M KCl, 30 mM MgCl_2_). Spin the cytoplasmic fraction at 15,000g for 15 min at 4 °C. Transfer the supernatant to a new tube and add 100% glycerol to a final concentration of 10%. Aliquot the cytoplasmic extracts and store at −80 °C before use.

To measure the mRNA stability in cytoplasmic extracts, 0.5 μg mRNA reporters are incubated in a mixture containing 1.5 μL of ATP (10 mM, NEB), 4 μL of 10% polyvinyl alcohol (Sigma-Aldrich), and 8 μL of cytoplasmic extracts at 30 °C for various times (0, 10, 30 min). Equal amount of firefly luciferase RNA is included as spike-in control followed by phenol: chloroform RNA precipitation. Purified RNA was dissolved in nuclease-free water for reverse transcription and PCR.

#### Real-time quantitative PCR

Purified total RNAs are reverse transcribed by High Capacity complementary DNA Reverse Transcription Kit (Invitrogen) using Random Primers (Thermo Fisher Scientific). Real-time PCR analysis is conducted using Power SYBR Green PCR Master Mix (Applied Biosystems) and carried on a Light Cycler 480 Real-Time PCR System (Roche Applied Science). Specific primers for amplifying each target genes are listed in Table S1.

#### cDNA library construction and deep sequencing

Purified total RNAs are suspended in nuclease-free water for reverse transcription.

In brief, RNA samples are mixed with 1 μl 10 mM dNTP and 2 pmol reverse primer overlap with 3’end of GFP (oligo 2) and incubated at 65 °C for 5 min, followed by incubation on ice for 5 min. The reverse transcription is carried out by incubating the reaction mixture with 1 × First-Strand Buffer, 10 mM DTT, 40 U RNaseOUT and 200 U SuperScript III at 50 °C for 60 min followed by heating at 70°C for 15minutes. First-strand cDNA is then used as the template for PCR amplification catalyzed by the Phusion High-Fidelity enzyme (NEB). The PCR is performed in a 20 μL reaction (1× HF buffer, 0.2 mM dNTP, 0.5 μM forward and reverse primers and 0.5 U Phusion polymerase) with barcoded primers (Table S1). PCR is set up based on the following condition: 16 cycles of 98 °C, 10 s; 60 °C, 20 s; 72 °C, 10 s followed by 72 °C, 10 min. The PCR products with the expected size 163 base pairs are excised from a 8% polyacrylamide TBE gel (Invitrogen). The DNA products are recovered from DNA gel elution buffer (300 mM NaCl, 1 mM EDTA) followed by quantification using Agilent BioAnalyzer DNA 1000 assay. Equal amounts of barcoded samples were pooled together followed by deep sequencing (Illumina HiSeq).

#### Lentiviral shRNAs and sgRNA

shRNA and sgRNA targeting sequences are designed based on RNAi consortium at Broad Institute (https://portals.broadinstitute.org/gpp/public/) and (http://crispr.mit.edu/) (Table S1). shRNA targeting sequences are cloned into DECIPHER pRSI9-U6-(sh)-UbiC-TagRFP-2A-Puro (Cellecta, CA), whereas sgRNA targeting sequences are cloned into LentiCRISPR v2 plasmid (Addgene plasmid #52961). Lentiviral particles are packaged using Lenti-X 293T cells (Clontech) according to the manufacturer’s instructions. Virus-containing supernatants are collected at 48 hr after transfection and filtered to eliminate cell contaminates. Cells are infected for 48 hr before selection by 2 mg/mL puromycin.

#### Immunofluorescence staining

HEK293-K^b^ cells are seeded in a glass bottom 24-well plate and grow overnight to ~ 70% confluence. Cells are transfected with Alexa Fluor-labeled RNA for 5 h by Lipofectamine MessengerMAX followed by fixation with 4% paraformaldehyde in PBS at room temperature for 10 min. The fixed cells are stained with 1 μM probe QUMA-1 for 30 min at 37 °C. Alternatively, cells are treated with 0.2% Triton X-100 at room temperature for 5 min for permeabilization followed by incubation with 1% BSA for 1 h. Cells are then incubated with anti-DCP2 antibody overnight at 4 °C and then with Alexa Fluor 594 goat anti-rabbit secondary antibody (Invitrogen, A-11007) for 1 h at room temperature. Hoechst 33342 (1:1,000 dilution)-stained compartments serve as markers of the nuclei. The glass slips are mounted on slides using nail polish and prepared for imaging using a Zeiss LSM710 confocal microscope.

#### Quantification and statistical analysis

##### Estimation of mRNA levels

The 3′ adapter CTCGAGCAGCTGGAAGATCG and low quality bases are trimmed by Cutadapt. The trimmed reads with length unequal to 10 nt are excluded. The count of each random sequence is calculated by a custom Perl script. A RPM value (reads per million) is obtained by dividing the resultant read count by total count.

##### Fluorescence intensity calculation

We sort transfected cells into 25D1 positive (25D1^H^), 25D1 negative (25D1^NEG^), GFP positive (GFP^H^) and GFP negative (GFP^NEG^) populations, followed by deep sequencing of the inserted 10 nt random sequences. For each random sequence, a 25D1 intensity is estimated as the average fluorescence intensity of 25D1^H^ and 25D1^NEG^, weighted by its RPM values in 25D1^H^ and 25D^NEG^ (Equation 1). The GFP intensity is estimated using the same method.

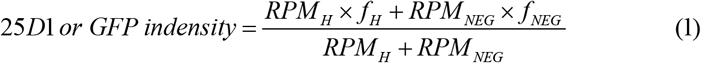

Where *RPM*_*H*_ and *RPM*_*NEG*_ are RPM values in 25D1 (GFP) positive and negative populations. *f*_*H*_ and *f*_*NEG*_ are mean fluorescence intensities in 25D1 (GFP) positive and negative populations.

##### Half-life estimation

We use a two-step method to estimate half-lives of random sequences without the guide of spike-in sequences. For each random sequence, a pre-half-life value is first estimated by Equation 2 and 3.

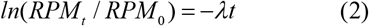

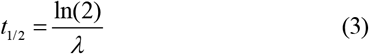

Where *RPM*_*t*_ and *RPM*_*0*_ are RPM values at time point *t* and 0. The slope λ is estimated by linear regression. Without correcting RPM value by spike-in sequences, estimated half-life values of the most stable sequences could be negative. Therefore, we normalize RPM values of each sequence by mean value of the most stable 1000 sequences. The half-life of each sequence (not including the most 1000 stable sequences) is re-estimated by Equation 2 and 3, using normalized RPM values. For linear regression in equation 2, only the sequences with estimated *P* value <0.05 and R^2^ >0.5 are used in downstream analysis.

## References

1. Bicknell, A.A. & Ricci, E.P. When mRNA translation meets decay. Biochem Soc Trans 45, 339–351 (2017).

2. Schwartz, D.C. & Parker, R. Mutations in translation initiation factors lead to increased rates of deadenylation and decapping of mRNAs in Saccharomyces cerevisiae. Mol Cell Biol 19, 5247–5256 (1999).

3. Kurosaki, T., Popp, M.W. & Maquat, L.E. Quality and quantity control of gene expression by nonsense-mediated mRNA decay. Nat Rev Mol Cell Biol 20, 406–420 (2019).

4. Presnyak, V. et al. Codon optimality is a major determinant of mRNA stability. Cell 160, 1111–1124 (2015).

5. Chan, L.Y., Mugler, C.F., Heinrich, S., Vallotton, P. & Weis, K. Non-invasive measurement of mRNA decay reveals translation initiation as the major determinant of mRNA stability. Elife 7 (2018).

6. Hinnebusch, A.G., Ivanov, I.P. & Sonenberg, N. Translational control by 5’-untranslated regions of eukaryotic mRNAs. Science 352, 1413–1416 (2016).

7. Dvir, S. et al. Deciphering the rules by which 5’-UTR sequences affect protein expression in yeast. Proc Natl Acad Sci U S A 110, E2792–2801 (2013).

8. Noderer, W.L. et al. Quantitative analysis of mammalian translation initiation sites by FACS-seq. Mol Syst Biol 10, 748 (2014).

9. Cuperus, J.T. et al. Deep learning of the regulatory grammar of yeast 5’ untranslated regions from 500,000 random sequences. Genome Res 27, 2015–2024 (2017).

10. Sample, P.J. et al. Human 5’ UTR design and variant effect prediction from a massively parallel translation assay. Nat Biotechnol 37, 803–809 (2019).

11. Starck, S.R. et al. Translation from the 5’ untranslated region shapes the integrated stress response. Science 351, aad3867 (2016).

12. Dersh, D., Yewdell, J.W. & Wei, J. A SIINFEKL-Based System to Measure MHC Class I Antigen Presentation Efficiency and Kinetics. Methods Mol Biol 1988, 109–122 (2019).

13. Ingolia, N.T., Lareau, L.F. & Weissman, J.S. Ribosome profiling of mouse embryonic stem cells reveals the complexity and dynamics of mammalian proteomes. Cell 147, 789–802 (2011).

14. Lee, S., Liu, B., Huang, S.X., Shen, B. & Qian, S.B. Global mapping of translation initiation sites in mammalian cells at single-nucleotide resolution. Proc Natl Acad Sci U S A 109, E2424–2432 (2012).

15. Kearse, M.G. & Wilusz, J.E. Non-AUG translation: a new start for protein synthesis in eukaryotes. Genes Dev 31, 1717–1731 (2017).

16. Fritz, D.T., Ford, L.P. & Wilusz, J. An in vitro assay to study regulated mRNA stability. Sci STKE 2000, pl1 (2000).

17. Kim, Y.K. & Maquat, L.E. UPFront and center in RNA decay: UPF1 in nonsense-mediated mRNA decay and beyond. RNA 25, 407–422 (2019).

18. Hogg, J.R. & Goff, S.P. Upf1 senses 3’UTR length to potentiate mRNA decay. Cell 143, 379–389 (2010).

19. Kwok, C.K., Marsico, G. & Balasubramanian, S. Detecting RNA G-Quadruplexes (rG4s) in the Transcriptome. Cold Spring Harb Perspect Biol 10 (2018).

20. Fay, M.M., Lyons, S.M. & Ivanov, P. RNA G-Quadruplexes in Biology: Principles and Molecular Mechanisms. J Mol Biol 429, 2127–2147 (2017).

21. Chen, X.C. et al. Tracking the Dynamic Folding and Unfolding of RNA G-Quadruplexes in Live Cells. Angew Chem Int Ed Engl 57, 4702–4706 (2018).

22. Kumari, S., Bugaut, A., Huppert, J.L. & Balasubramanian, S. An RNA G-quadruplex in the 5’ UTR of the NRAS proto-oncogene modulates translation. Nat Chem Biol 3, 218–221 (2007).

23. Herdy, B. et al. Analysis of NRAS RNA G-quadruplex binding proteins reveals DDX3X as a novel interactor of cellular G-quadruplex containing transcripts. Nucleic Acids Res 46, 11592–11604 (2018).

24. Weingarten-Gabbay, S. et al. Comparative genetics. Systematic discovery of cap-independent translation sequences in human and viral genomes. Science 351 (2016).

25. Wahle, E. & Winkler, G.S. RNA decay machines: deadenylation by the Ccr4-not and Pan2-Pan3 complexes. Biochim Biophys Acta 1829, 561–570 (2013).

26. Gilbert, W.V., Zhou, K., Butler, T.K. & Doudna, J.A. Cap-independent translation is required for starvation-induced differentiation in yeast. Science 317, 1224–1227 (2007).

27. Mayr, C. Regulation by 3’-Untranslated Regions. Annu Rev Genet 51, 171–194 (2017).

## References

1. D. T. Fritz, L. P. Ford, J. Wilusz, An in vitro assay to study regulated mRNA stability. Sci STKE 2000, pl1 (2000).

